# Metagenomic coverage bias at transcription start sites is correlated with gene expression

**DOI:** 10.1101/2024.05.09.593333

**Authors:** Gordon Qian, Izaak Coleman, Tal Korem, Joshua W. K. Ho

**Affiliations:** School of Biomedical Sciences, Li Ka Shing Faculty of Medicine, The University of Hong Kong, Pokfulam, Hong Kong SAR; Laboratory of Data Discovery for Health (D^2^4H), Hong Kong Science Park, Hong Kong SAR; Program for Mathematical Genomics, Department of Systems Biology, Columbia University Irving Medical Center, New York, NY; Department of Obstetrics and Gynecology, Columbia University Irving Medical Center, New York, NY

**Keywords:** Metagenomics, Gut microbiome, Next-generation sequencing

## Abstract

Metagenomic sequencing is presumed to provide unbiased sampling of all the genetic material in a sample. Downstream analysis methods, such as binning, gene copy number analysis, structural variations, or single nucleotide polymorphism analysis, commonly assume an even distribution across the genome after accounting for known artefacts such as GC content. We discovered coverage bias across gut microbiome species, manifesting as a difference in coverage before and after bacterial transcription start sites. Using matched metatranscriptomic and metagenomic sequencing data, we demonstrate that this bias correlates with gene expression. Potential artefacts such as the sequencing technology, reference genome used for alignment, and mappability bias were investigated across multiple datasets and shown to not be factors for association. While GC bias was found correlated with coverage bias, the association of coverage bias with gene expression remains significant after adjusting for GC bias. Paired-end read mapping demonstrated an enrichment in 5’ read ends immediately downstream of the TSS which was partly a byproduct of unmapped reads upstream of the TSS. Our observations suggest the existence of strain-level variation where sequence variation in the promoter site region is preventing proper read alignment to the reference genome. The correlation of this phenomenon with gene expression may also reflect evolutionary footprints for fine-tuning the regulation of gene expression. Understanding the source of this sequence variation and the biological implications of this artefact will be useful not only to better characterise microbial functions but also to improve interpretations of strain level dynamics.

**Importance:** Sequencing coverage calculated from metagenomic sequencing data is extensively used in the microbiome field, providing valuable information about microbial abundances, gene (functional) abundances, growth rates, and genomic variations. Understanding factors that impact the distribution of coverage along genomes is therefore important for multiple applications. In this study, we report on uneven read coverage across the transcription start sites of bacterial genomes that is correlated with gene expression levels. We determine that this bias is independent of multiple factors including GC bias, and arises due to higher strain divergence from reference genomes upstream of the transcript start site. We propose that evolutionary finetuning of gene expression in competitive microbial ecosystems can drive genetic mutations at the promoter site. Our findings suggest the potential to glean gene regulatory information from metagenomic data, and better understand how ecological factors shape genomes in the microbiome and their sequencing coverage.

## Introduction

Metagenomic sequencing has allowed studies to profile all genomic content in a sample at a massively parallel and high-throughput manner. As such, it is also important to understand the artefacts it may contain. An important type of artefact is read coverage bias. For example, GC content is known to drive under- and over-representation of genomic sites within a genome (1), and was demonstrated to drive non-random fragmentation, influencing fragment counts and generating non-uniform coverage, with fragments more likely to start within a CpG dinucleotide (2, 3). PCR amplification bias has been demonstrated to be a major source for coverage bias, with amplicons of GC > 65% diminished by a hundred-fold compared to the mid-GC reference loci, while those with GC < 12% are diminished by ten-fold (1, 4). Methods for GC-correction have been developed (3), although they generally rely on a fragmentation model which may not always accurately represent all samples or microbial compositions (3). Yet another source of coverage bias arises from read mappability, which is caused by lack of uniqueness of the mapped read (5, 6). In cases where k-mer sequences are found perfectly repeated on the reference, alignment algorithms will struggle to correctly assign read positions, leading to underrepresentation of coverage in areas of low complexity (7).

In addition to the existence of technical artefacts that influence coverage bias, biological phenomena have also been identified. In eukaryotic samples, the presence of chromatin and nucleosomes can influence the sampling of DNA sequences, because they protect the DNA from fragmentation or enzymatic digestion (8). A recent study has exploited this fact and demonstrated read coverage of circulating cell-free DNA in the blood correlates with gene expression due to different fragmentation profiles of gene regions from altered chromatin methylation (9). This observation was then utilised to classify cancer types due to differences in their gene expression profiles from normal cells. Despite the absence of chromatin and nucleosome structures in prokaryotes, alternative mechanisms of DNA coiling or protein binding may potentially exhibit similar effects on read coverage. This includes phenomena such as supercoiling, which has been implicated in bacterial gene expression (10), and large structural rearrangements that they cause, termed plectonemes (11). It was demonstrated that plectonemes are commonly found upstream of gene transcription promoter sites, possibly indicating a role in gene regulation similar to nucleosomes in eukaryotes (11).

Metagenomics has proved extremely effective in capturing a comprehensive snapshot of microbial genomes. This has facilitated significant advancements in our capabilities for high-throughput analysis of complex microbial communities and our understanding of previously unculturable microbes and their role in human health (12, 13). Therefore, understanding properties, drivers and biases of read coverage bias becomes critical for applications such as binning, gene copy number analysis, structural variant calling and SNP calling (14–20), which all rely on the assumption that sequencing coverage across genomes is more or less even.

In this work, we thoroughly investigate coverage patterns across genomes from human gut metagenomic samples to assess the extent of coverage bias and identify potential technical or biological sources. We specifically look at read coverage at transcription start sites, with the initial hypothesis that biological mechanisms related to gene expression such as polymerase binding or supercoiling influence coverage. We identify systematic coverage bias enriched at TSSs over various datasets and species genomes, and show that this correlates with gene expression in a matched dataset of metagenomic and metatranscriptomic sequencing. To investigate potential explanations for this, we rule out the possibility of off-target RNA sequencing and instead discover an enrichment of unmapped reads upstream of the TSS. We demonstrate this arises due to increased sequence heterogeneity compared to reference genomes used for alignment, which we propose to be the result of competitive selective pressures within the microbiome.

## Methods

### Metagenome sequencing data

Matched metagenomic and metatranscriptomic data of gut stool samples were downloaded from NCBI BioProject PRJNA188481. Analysis to show correlation of TSS coverage bias across samples and datasets utilised 2 additional gut metagenome cohort studies, including a hypertensive cohort (22) and a cohort looking at antibiotic drugs on antibiotic resistance in the GM (23). Oxford Nanopore and PacBio Hi-Fi long-read metagenomic stool datasets utilised for comparison of TSS coverage bias patterns across sequencing platforms were downloaded from PRJNA820119 (25) and PRJNA754443 (24), respectively. Matched sample Illumina sequencing was also downloaded for ten samples (CD91 - CD100) from the Oxford Nanopore dataset, which was used to assess coverage bias at TSSs when aligning short-reads to long-read references. As a control, a matched long-read/short-read bacterial mock community 20 Strain Staggered Mix Genomic Material (ATCC MSA-1003) was downloaded from the following accessions: SRR11606871 (PacBio HiFi) and SRR8359173 (Illumina) (28).

### Computing metagenomic coverage and metatranscriptomic quantification

Short-read Illumina metagenomic sequencing samples were directly aligned to reference genomes with Bowtie2 using default parameters (38). For PacBio Hi-Fi and Oxford Nanopore long-reads, bwa-mem was used for alignment with the appropriate parameters, ‘-x pacbio’ and ‘-x ont2d’, respectively.

Reference genome gene annotations were called using Prodigal v2.6.3 with default parameters (29). Gene annotation lists were filtered based on the rule that the gene start must be at least more than 500 bps away from their neighbouring gene starts. This was to isolate coverage bias effects to individual TSSs, and prevent possible coverage bias phenomenons at TSSs affecting the coverage values of neighbouring TSS windows. Coverage of alignments were retrieved with bamCoverage from the deepTools bioinformatic suite using default parameters (39). Metatranscriptomic sequencing reads were aligned to gene reference sequences using kallisto quant (40). Gene expression is quantified under the normalisation of transcripts per million (TPM). We then further categorised genes into 4 expression groups, using the following thresholds. Zero: TPM ==0, Low: TPM > 0 & TPM < 100, Mid: TPM >= 100 & TPM < 1000, High: TPM >= 1000.

### Coverage and GC bias calculation

Coverage across a genome of length L can be defined by a series of read counts at each position, 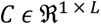, where *c_i_* defines the read count at the i^th^ position in the genome. Along a genome, we define a set of N TSS locations, 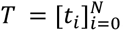, where *t_i_* ∈ [0, *L*]. The coverage bias of a TSS is defined by considering a window, w, of 1001 coverage values, 500 bps upstream and downstream of the TSS loci, 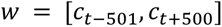, and standard scaling all values within this window, 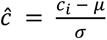, where μ and σ represent the mean and variance of w. Coverage bias, *β_cv_*, is calculated by taking the mean difference of the downstream scaled coverage and upstream scaled coverage, 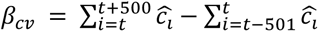. GC content bias at the TSS was simply calculated by the difference in GC% downstream of the TSS to GC% upstream of the TSS.

### Metagenome read simulation

For read mappability bias testing, paired-end Illumina metagenomic sequencing reads were simulated from reference genomes using art_illumina from the ART suite (26). The following parameters were used for simulation, -na -ss HS25 -p -l 150 -c 500000 -m200 -s 10.

### Systematic coverage bias analysis

The analysis of coverage bias at the TSS and its correlation with gene expression was performed systematically across all 8 samples from the BioProject PRJNA188481. Species abundance was estimated using MetaPhlAn3 (41). Species considered for analysis include those present at a relative abundance of at least 1% in at least 5 samples. For each selected species, up to 3 random genome assemblies were selected for alignment where the average result of all genome assemblies is taken for a species. Genome assemblies were sourced from Refseq with the criteria of having an assembly level of “Complete Genome”. This captured 7 unique species with a total of 20 genome assemblies:

| Taxonomic ID | Taxonomic Name | Refseq Accession and assembly |
| --- | --- | --- |
| 820 | *Bacteroides uniformis* | GCF_006742345.1_ASM674234v1<br>GCF_018289175.1_ASM1828917v1<br>GCF_018289375.1_ASM1828937v1 |
| 821 | *Phocaeicola vulgatus* | GCF_000012825.1_ASM1282v1<br>GCF_008728395.1_ASM872839v1<br>GCF_018289355.1_ASM1828935v1 |
| 28117 | *Alistipes putredinis* | GCF_000154465.1_ASM15446v1<br>GCF_015549455.1_ASM1554945v1<br>GCF_015554085.1_ASM1555408v1 |
| 823 | *Parabacteroides distasonis* | GCF_000012845.1_ASM1284v1<br>GCF_006149185.1_ASM614918v1<br>GCF_006739545.1_ASM673954v1 |
| 39485 | *Lachnospira eligens* | GCF_000146185.1_ASM14618v1<br>GCF_020735745.1_ASM2073574v1 |
| 1150298 | *Fusicatenibacter saccharivorans* | GCF_001404555.1_14207_7_46<br>GCF_001405555.1_13414_6_33<br>GCF_001406335.1_14207_7_8 |
| 853 | *Faecalibacterium prausnitzii* | GCF_002586945.1_ASM258694v1<br>GCF_002734145.1_ASM273414v1<br>GCF_003293635.1_ASM329363v1 |

Gene expression (TPM) was modelled with a mixed model of fixed and random effects using the LmerTest T library. Fixed effects included coverage bias and GC bias. Random effects included the metagenomic sample, and the microbial species. TPM was log-transformed with the addition of a small pseudocount, and then standard scaled along with coverage bias and GC bias. TSSs with no coverage across their 1kb window were excluded from the model.

### Permutation significance testing of bias measurements

To assess the statistical significance of the average bias metric across all TSSs along a genome, we compared it to a distribution bias values from randomised TSS locations along the genome as a negative control. For a reference genome G, with N number of annotated TSSs, N random positions were randomly selected along the genome such that the distances between them maintains the same distances that of the original TSSs. The average read bias value was measured across all N positions, and this was repeated 500 times to form the null distribution. Significance was evaluated using a 1- tailed hypothesis test. This permutation test was similarly performed to measure GC bias significance.

### Bias correlation analysis

The robustness of TSS read bias across samples was estimated through pairwise Pearson correlation coefficients by comparing the bias vectors of each sample. This was performed systematically across 4 datasets and 38 samples. The average Pearson R value was calculated for all pairwise permutations within-dataset and across-dataset. Species considered in this analysis were filtered under the requirement that it was present at a relative abundance of 1% in at least 5 samples in each dataset. This captured 5 species, although only 4 were included in analysis as reference genomes for *[Eubacterium] rectale* was not found in Refseq. Bias metrics also seemed to deviate depending on the strain reference genome used, and therefore where there existed more than 1 complete reference genome for a species, up to 3 different strain references were used for alignment. Correlation scores across strains were averaged across strains of the same species and these values were used for independent t-test statistical comparison of samples across datasets. Samples where the species was not present in at least 1% abundance was not considered in the average correlation score. The resulting reference genome assemblies used in this analysis:

| Taxonomic ID | Taxonomic Name | Refseq Accession and assembly |
| --- | --- | --- |
| 820 | *Bacteroides uniformis* | GCF_006742345.1_ASM674234v1<br>GCF_018289175.1_ASM1828917v1<br>GCF_018289375.1_ASM1828937v1 |
| 821 | *Phocaeicola vulgatus* | GCF_000012825.1_ASM1282v1<br>GCF_008728395.1_ASM872839v1<br>GCF_018289355.1_ASM1828935v1 |
| 28117 | *Alistipes putredinis* | GCF_000154465.1_ASM15446v1<br>GCF_015549455.1_ASM1554945v1<br>GCF_015554085.1_ASM1555408v1 |
| 853 | *Faecalibacterium prausnitzii* | GCF_002586945.1_ASM258694v1<br>GCF_002734145.1_ASM273414v1<br>GCF_003293635.1_ASM329363v1 |

For assessing the robustness of read bias across samples from different sequencing platforms, species with at least 0.5% abundance in at least 4 samples for each dataset were considered. This resulted in 4 species:

| Taxonomic ID | Taxonomic Name | Refseq Accession and assembly |
| --- | --- | --- |
| 853 | *Faecalibacterium prausnitzii* | GCF_002586945.1_ASM258694v1<br>GCF_002734145.1_ASM273414v1<br>GCF_003293635.1_ASM329363v1 |
| 74426 | *Collinsella aerofaciens* | GCF_002736145.1_ASM273614v1<br>GCF_003856815.1_ASM385681v1<br>GCF_005576555.1_ASM557655v1 |
| 166486 | *Roseburia intestinalis* | GCF_025151715.1_ASM2515171v1 |
| 649756 | *Anaerostipes hadrus* | GCF_001998765.1_ASM199876v1<br>GCF_902388025.1_UHGG_MGYG-HGUT-02528 |

### Paired-end 5’ read enrichment analysis

5’ read enrichment immediately downstream of the TSS was calculated by considering a 50 bp window starting from 10 bps upstream of the TSS, and taking the ratio of 5’ read to 3’ reads in this window. To investigate whether observations were unique to the TSS, we repeated this analysis with an added “offset” to the window, where an offset of 100 would shift the original 50 bp windows downstream the gene by 100 bps. Reads were counted under two frameworks, which we refer to as paired and single mapped. Paired counting only considers reads where both the reads in each pair were successfully aligned with Bowtie2, whereas single mapped counting also included reads whose pair was not aligned. Genes shorter than 500bp were discarded from this analysis so that an offset of 400 bp does not extend past the end of the gene.

### Reference genome and long read reference analysis

Read alignment to long-reads followed the same method as to reference genomes mentioned above, except with the additional -a parameter in Bowtie2 to allow for multi-mapping, as long-reads can exist as overlapping regions. Long-reads shorter than 10,000 bp were removed from analysis to minimise segmented gene regions at long read borders, and subsampled to 100,000 sequences prior to indexing and alignment due to significant computational speed up. Prodigal was run on long-reads to annotate gene regions. The gene sequences were annotated by mapping to genes annotated on reference genomes using minimap2. This allows us to match long reads to their respective reference genome and annotate them similarly. Additionally, any annotated genes within 500 bps of long read ends were filtered out in order to have sufficient sequence coverage information to calculate bias.

### Strain diversity read alignment analysis

This analysis relies on the idea that singleton reads arise from misalignment of its pair due to discrepancies to the reference genome, specifically at the intergenic region. For this reason, it would be advantageous to test this hypothesis with sequencing libraries with larger fragment length distributions. This allows singleton reads aligned at the beginning of a gene, in the reverse orientation to the gene sense, to have their read pair likely exist upstream in the intergenic region. This is in contrast to short fragment length libraries where both read pairs may share a considerable amount of overlap. We used 10 samples from the antibiotic resistance gene (ARG) metagenomic study for this analysis due to its larger fragment lengths of ∼ 300 bps (23). Species with at least 1% abundance in at least 5 samples were examined in this analysis. This captured 8 species with a total of 24 reference genomes:

| Taxonomic ID | Taxonomic Name | Refseq Accession and assembly |
| --- | --- | --- |
| 310297 | *Phocaeicola plebeius* | GCF_000187895.1_ASM18789v1<br>GCF_003437535.1_ASM343753v1<br>GCF_003438135.1_ASM343813v1 |
| 820 | *Bacteroides uniformis* | GCF_006742345.1_ASM674234v1<br>GCF_018289175.1_ASM1828917v1<br>GCF_018289375.1_ASM1828937v1 |
| 821 | *Phocaeicola vulgatus* | GCF_000012825.1_ASM1282v1<br>GCF_008728395.1_ASM872839v1<br>GCF_018289355.1_ASM1828935v1 |
| 28117 | *Alistipes putredinis* | GCF_000154465.1_ASM15446v1<br>GCF_015549455.1_ASM1554945v1<br>GCF_015554085.1_ASM1555408v1 |
| 823 | *Parabacteroides distasonis* | GCF_000012845.1_ASM1284v1<br>GCF_006149185.1_ASM614918v1<br>GCF_006739545.1_ASM673954v1 |
| 301302 | *Roseburia faecis* | GCF_001405615.1_13414_6_47<br>GCF_001406815.1_M72<br>GCF_009718405.1_ASM971840v1 |
| 40518 | *Ruminococcus bromii* | GCF_002834165.1_ASM283416v1<br>GCF_002834225.1_ASM283422v1<br>GCF_002834235.1_ASM283423v1 |
| 853 | *Faecalibacterium prausnitzii* | GCF_002586945.1_ASM258694v1<br>GCF_002734145.1_ASM273414v1<br>GCF_003293635.1_ASM329363v1 |

A similar approach to pair-end enrichment analysis was used where we defined a window, *w*, immediately downstream of the TSS and captured all, *n*, 3’-end reads (both singleton reads and pair aligned reads). The 5’ pair of these reads will theoretically be located upstream of the TSS, regardless of if they were actually successfully mapped by Bowtie2. These reads are then aligned back to a reference proximal region of the TSS, *r*, which we define as a broad region 300 bps upstream the TSS to 200 bps downstream the TSS. This was performed by semi-global alignment (Pederson, 2016), global alignment with respect to the shorter sequence i.e. the read. The mean alignment score of all reads is used to present the heterogeneity of the gene’s upstream intergenic region, where a low average alignment score would reflect high heterogeneity due to differences from the reference genome and sample-specific strain genomes. The negative control was performed for each TSS by randomly selecting *n* number of reads and calculating the average alignment score to the TSS reference sequence, *r*. This reflects the distribution of alignment scores for real random alignments. The positive control, demonstrating alignment scores of reads in the intragenic region, was performed by shifting the target window of interest, *w*, downstream into the gene body by *f* bps, where *f* is the average fragment length. This ensures 3’-end reads captured have paired reads within the genic region, which should not contain the degree of heterogeneity as intergenic regions, according to our hypothesis.

## Results

### Uneven metagenomic coverage around bacterial transcription start sites

To investigate coverage at the gene transcription start site (TSS), we aligned stool metagenomic sequencing reads to species representative reference genomes and quantified coverage around all TSSs per genome (**Fig. 1a**). This was performed across all unique metagenomic samples (N=8) from the Franzosa et al. dataset (21), considering all species with at least 1% abundance in at least 5 samples. By visualising the coverage 500 bps downstream and upstream of all TSSs in the genome, we observed non-uniform coverage either downstream or upstream of the TSS (**Fig. 1b**). To quantify this observation, we calculated the difference between the coverage 500 bp upstream and downstream of the TSS, henceforth termed “TSS coverage bias”, across all TSSs. We found that the average TSS coverage was significantly higher than a null distribution in which coverage is unrelated to TSSs across 50 of 52 species-sample combinations (Methods; Permutation *p*<0.05; **Fig. 1c**, an example of *Faecalibacterium prausnitzii* in one sample is in **Fig. 1d**). This demonstrates that the specific positioning of TSSs along the genome is associated with observed coverage bias, where on average a greater number of reads are mapped downstream of the TSS.

**Figure 1.**
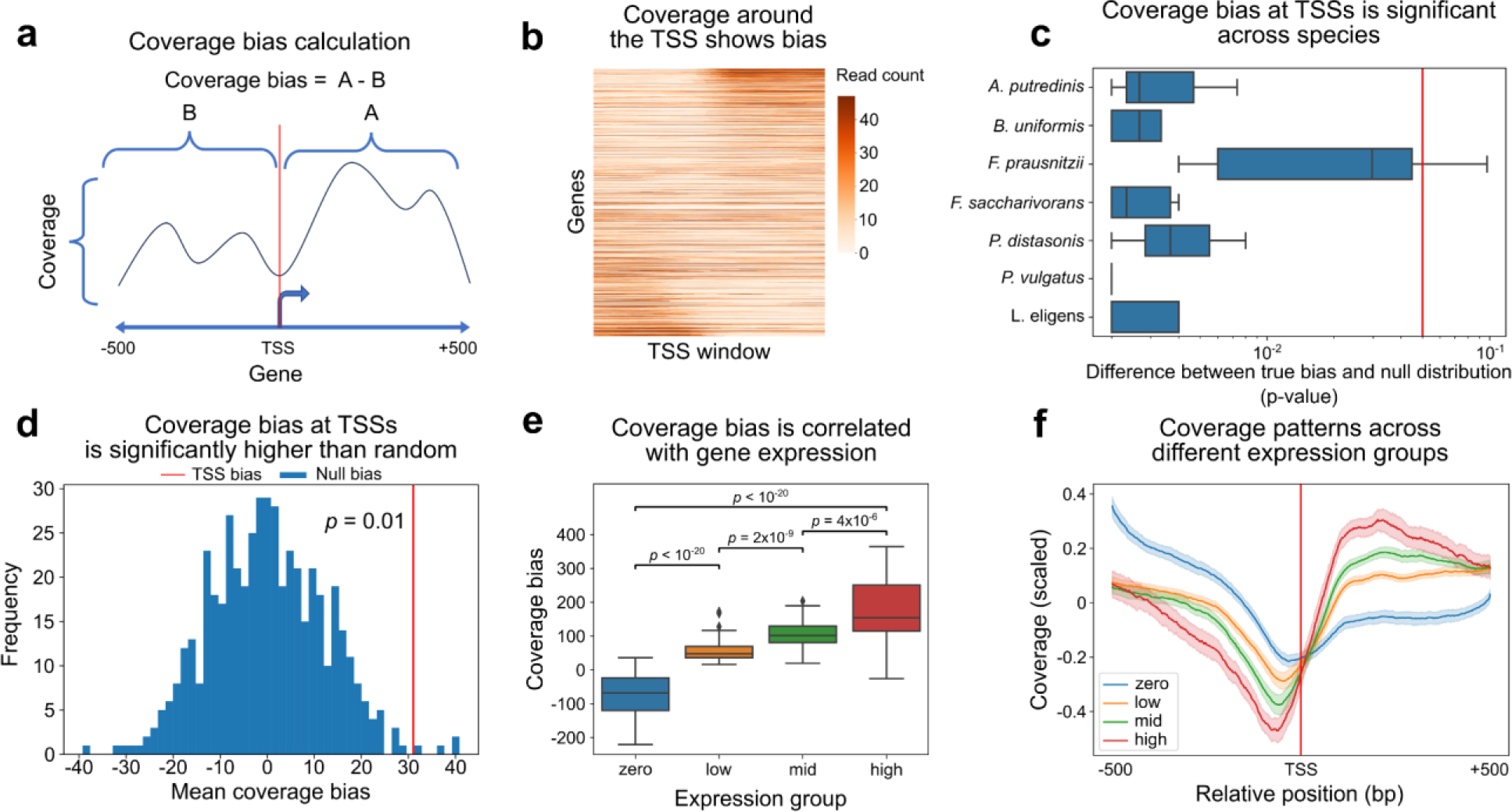
Uneven coverage across transcription start sites (TSS) is correlated with gene expression. **(a)** Diagram illustrating coverage bias calculation. **(b)** Heatmap of raw short-read coverage from one sample across gene transcription start sites. Shown are all windows 500bps upstream and 500bps downstream of all TSSs in the reference genome of *F. prausnitzii*. Genes are ordered by coverage bias. **(c)** Boxplot (line, median; box, IQR; whiskers, 1.5xIQR) of permutation p values of coverage bias enrichment in 8 samples from Franzosa’s dataset compared to a null distribution of random TSS placement, across various species genomes. The vertical red line indicates the threshold for significance of p = 0.05. **(d)** An example histogram of the average coverage bias distribution from 1000 random null permutations of TSS positions for *F. prausnitzii* in one sample. The vertical red line indicates the average coverage bias at the true TSSs in the same sample. **(e)** Boxplot of average coverage bias in different gene expression groups across different samples and species genomes. P - paired t-test. **(f)** Coverage in the 1 kbp around the TSS across all genes for 7 species from 8 samples from the Franzosa dataset (line, average; area, 95% confidence interval). coverage is scaled (Methods) and grouped by gene expression.

### Coverage bias is correlated with gene expression

We next investigated whether the coverage bias at a specific TSS is associated with the expression levels of the respective gene. To this end, we quantified the gene expression of all samples using its matched metatranscriptomic sequencing data and categorised the expression level (transcripts per million, TPM) of each gene as zero, low, mid, and high (Methods). The coverage bias of all genes within an expression group was then averaged for each sample per species. We found that coverage bias increased with gene expression across species (Kruskal p < 10^-20^, **Fig. 1e,f**). This positive correlation also remained true when stratifying by individual species (**Suppl. Fig. 1**). Average coverage across the TSS region dips approximately 100 bps upstream of the TSSs and peaks approximately 100 bps downstream of the TSSs. This pattern is consistent across all species-stratified analysis (**Suppl. Fig. 2**).

### TSS coverage bias is consistent across datasets

In order to determine if coverage bias exists as a systemic or sample-specific phenomenon, we evaluated the similarity of TSS coverage bias across all abundant taxa (at least 1% abundance in at least 5 samples of each dataset) from three different datasets: 10 samples from a cohort studying gut microbiome and hypertension (HYP) (22), 10 samples from a cohort studying antibiotic resistance genes in the GM (ARG) (23) and the 8 samples from Franzosa et al. (Franzosa). As these samples were sequenced by different labs, high similarity across samples would reduce the chances of the read bias signal being caused by protocol-specific technical artefacts. We found that the average pairwise correlation of TSS coverage bias between samples from the same dataset (Pearson R=0.34±0.034) was similar to the correlation of samples across datasets (Pearson R=0.30±0.024) (t-test p=0.19 for difference between inter- and intra- correlations; **Fig. 2a**). As expected, a null model of random read positions had no correlation across samples (Pearson R=0.00029±0.0014; **Suppl. Fig. 3a**). Altogether, our results indicate that coverage bias around the TSS is a generalizable phenomenon consistent across several datasets.

**Figure 2.**
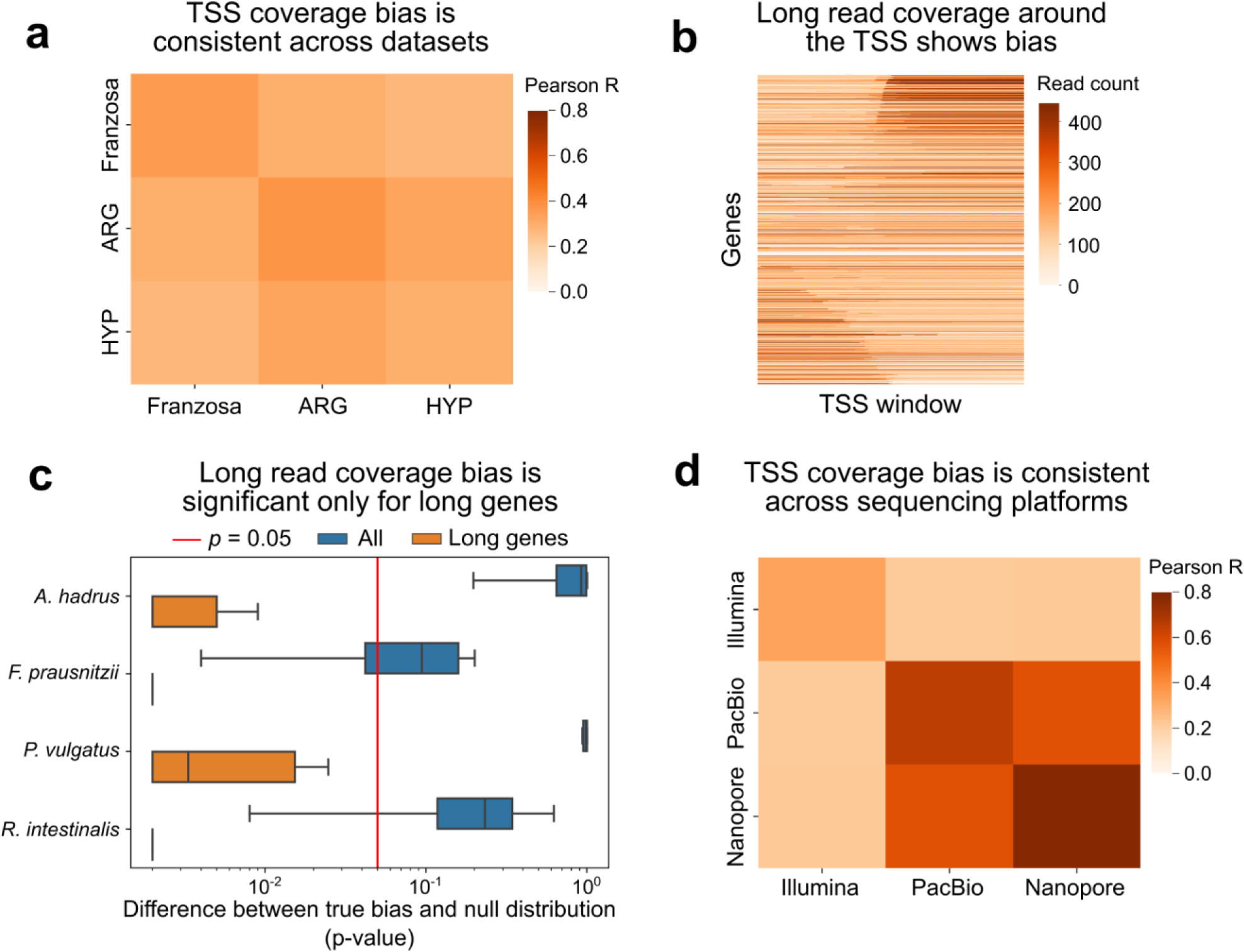
Coverage bias is reproducible across datasets. **(a)** Heatmap of Pearson correlation scores between coverage bias values across all TSSs in samples from different datasets. Pearson correlation scores were calculated as the average pairwise correlation for all sample comparisons per species. A total of 8 samples and 4 species (3 reference genome each) were assessed where average correlation scores for each reference genome were averaged for each species. **(b)** Heatmap of raw read coverage of PacBio long reads from one sample in the 500bps upstream and downstream of all TSSs in the reference genome of *F. prausnitzii*. **(c)** P-values of coverage bias enrichment in comparison to random null distributions using long reads from 16 samples across various species genomes, stratified by gene length. The vertical red line indicates p = 0.05. **(d)** Same as (a), for comparison between different samples sequenced using different sequencing platforms.

We also sought to identify whether this coverage bias was specific to Illumina short-read sequencing. To validate its presence across species and samples, we tested whether the average coverage bias at TSSs was higher than a null distribution across 6 PacBio long read HiFi sequenced samples (24) and 10 Oxford Nanopore long read sequenced gut metagenomic samples (25). To capture more shared species across platforms, we aligned reads to all species present at >0.5% in at least 4 samples in each dataset (Pacbio, Nanopore and Illumina). We found, unlike Illumina samples, only 6 of 56 species-sample combinations had a significantly higher TSS coverage bias than a random null **(Fig. 2b,c)**. We posit that this may be a result of long reads typically spanning multiple genic and intergenic regions, and hypothesised that the effect would be diminished when examining longer genes. Indeed, when considering only genes longer than 1 kbps and found that coverage bias enrichment was significant for 55 of 56 species-sample combinations (permutation *p* < 0.05; **Fig. 2c)**. This indicates coverage bias at the TSS is not a phenomenon specific to Illumina short-read sequencing.

We also compared the similarity of coverage bias between this short-read and long-read data. We found extremely high concordance between coverage bias for samples within the same dataset for long-read sequencing datasets (PacBio HiFi R=0.65±0.052; Oxford Nanopore R=0.78±0.02; R=mean±std of species means, **Fig. 2d**). Coverage bias correlation between PacBio HiFi samples and Oxford Nanopore samples also presented high concordance (R=0.57±0.037) whereas their correlation with Illumina sequenced samples was much lower (Illumina vs. Pacbio HiFi R=0.22±0.094, Illumina vs.

Oxford Nanopore R=0.22±0.12, **Fig 2d**). This likely arises due to the extended coverage of long-reads where the duality of either upstream or downstream alignment will cover the entire 500bp window whereas short reads do not, thus amplifying the bias signal. Regardless of the lower correlation score between short-read and long-read samples, this score is still significantly higher than the null model of randomised TSS positions (Illumina vs. PacBio HiFi p=0.0072, Illumina vs. Oxford Nanopore p=0.018, **Suppl. Fig. 3b**).

### TSS coverage bias is associated with GC-bias but not with mappability

We next sought to identify potential sources for the TSS coverage bias. We first examined if the observed coverage bias is caused by read mappability, i.e., a reduction of coverage caused by issues in mapping to low complexity or repetitive regions in the genome (5). To this end, we simulated reads uniformly from reference genomes (26) to reflect read sequencing from a sample (**Methods**) and re-aligned them back to the reference genome, such that the presence of any coverage bias at the TSSs would suggest bias due to mappability (**Fig. 3a**). The coverage bias observed after aligning these simulated reads to the TSS regions was similar to randomly selected regions across all reference genomes (permutation *p* > 0.05 for all 7 species; **Fig. 3b**). We also tested the similarity of coverage bias at TSSs from real sample reads with their simulated counterparts and found no correlation between them across all samples and selected species (**Fig. 3c**). This indicates mappability bias is not a major contributor to the TSS coverage bias that we observed.

**Figure 3.**
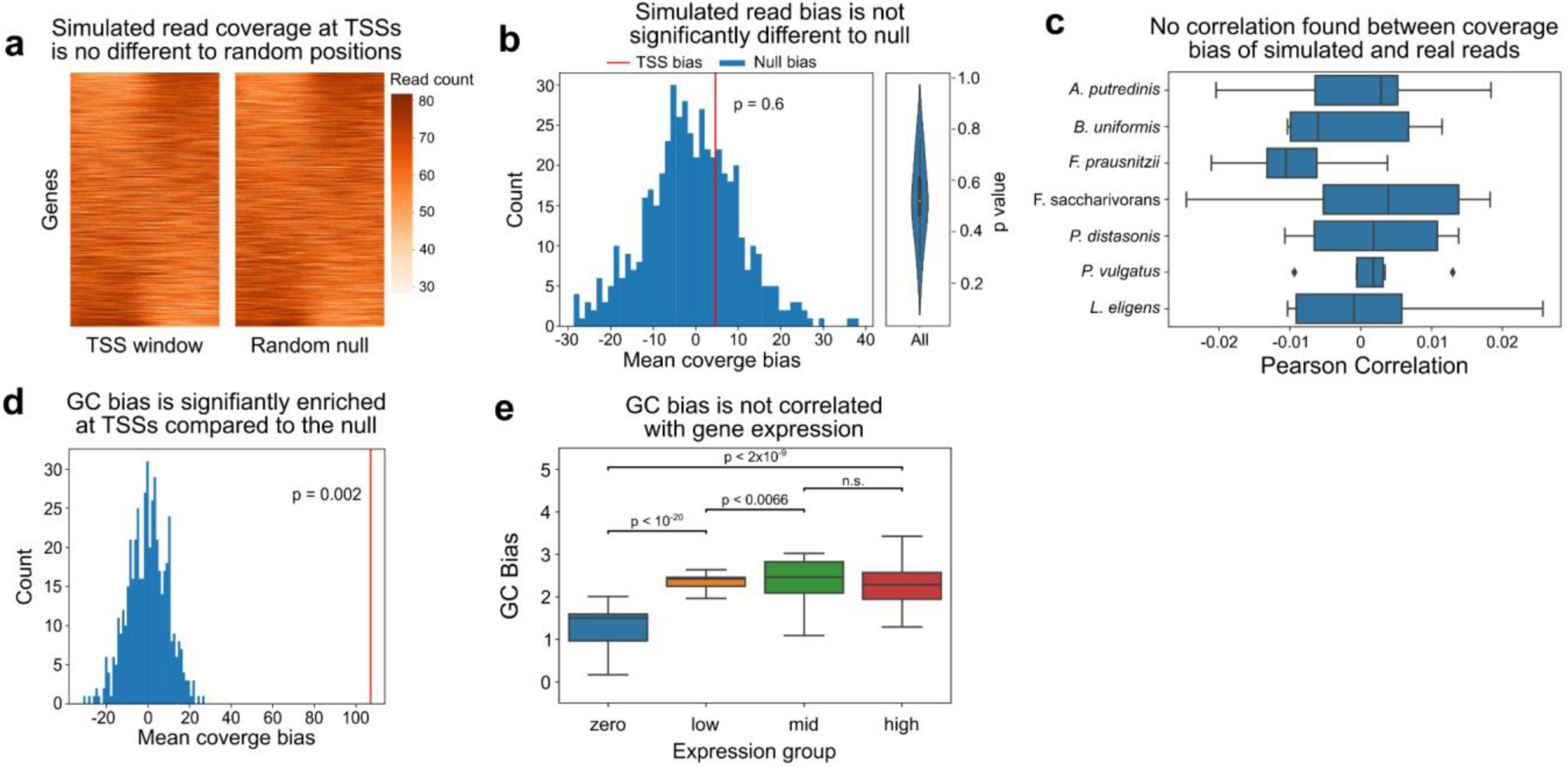
GC bias, but not read mappability, are associated with TSS coverage bias. **(a)** Heatmap of showing coverage of simulated reads in a 1kbp window surrounding the TSS (left) or randomly selected positions (right) on the *F. prausnitzii* reference genome. **(b)** Left, histogram of the average coverage biases in the 1kbp region around 1000 random genomic positions. The vertical red line indicates the average TSS coverage bias of simulated reads. Right, violin plot of permutation p-values for all reference genomes, demonstrating that simulated read coverage bias at TSSs is not significantly different than from random positions. **(c)** Boxplot (line, median; box, IQR; whiskers, 1.5xIQR) of Pearson correlation between coverage biases across TSSs from simulated reads and real metagenomic sample reads, across 8 samples. **(d)** Histogram of the average GC bias distribution from 1000 random null permutations of genomic positions. The vertical red line indicates the average GC bias at TSSs. **(e)** Boxplot of average GC bias for different gene expression groups across different samples and species genomes. P - paired t-test.

The GC content of sequences is known to influence sequencing coverage (27). To test the influence of GC content on TSS coverage bias, we calculated a GC bias metric similarly to coverage bias by taking the difference between the GC content downstream and upstream of the TSS (**Methods**). In all species genomes from the 8 Franzosa dataset samples, the mean GC bias across all TSSs was significantly higher in comparison to a null distribution of mean GC bias of randomly selected positions (permutation *p* = 0.002 for all species, **Fig. 3d** shows an example of *F. prausnitzii* reference genome). TSS GC bias is therefore associated with coverage bias and, therefore, if GC bias is associated with gene expression it could confound the association of coverage bias with gene expression. To test this, we evaluated the association of GC bias with gene expression in the same samples and species from the Franzosa dataset. Across all species (N = 7), the GC bias of the low, mid and high gene expression groups was higher than in the zero gene expression group (*p* < 10^-20^, mean GC bias of 1.21, 2.21, 2.38, 2.29 for zero, low, mid and high, respectively, **Fig. 3e**). However there was no ordinal relationship between GC bias in the low to high gene expression groups where the high gene expression group had a non-significant decrease in GC mean compared to the mid gene group (**Fig. 3e**). Finally, to systematically evaluate the total effect of different factors in the associations with coverage bias across samples and species, we fitted a linear mixed model using samples and species as random effects and TSS coverage bias and GC bias as fixed effects to predict gene expression. The resulting model demonstrated that both coverage bias and GC content were significantly associated with gene expression, with evidence for a stronger association with coverage bias (**Table 1**).

**Table 1.** Gene expression is associated with coverage bias even when correcting for GC bias.

| Linear Mixed Model | Estimate | Standard Error | t value | P > \|z\| |
| --- | --- | --- | --- | --- |
| Coverage bias | $9.672 \times 10^{-2}$ | $1.934 \times 10^{-3}$ | 50.01 | $< 2 \times 10^{-16}$ |
| GC bias | $4.218 \times 10^{-2}$ | $1.941 \times 10^{-3}$ | 21.73 | $< 2 \times 10^{-16}$ |

Gene expression was modelled under a linear mixed model with fixed effects for TSS coverage and gc bias for each individual gene. Random effects, sample and species, allowed grouping of observations. Gene expression (TPM) values, as well as bias variables, were log-transformed, scaled and centered. There were a total of 245,861 observations, 20 species groups and 8 sample groups. Model convergence by Restricted Maximum Likelihood (REML).

### The correlation of coverage bias and gene expression is not an artefact of RNA sequencing

The association between TSS coverage bias and gene expression could potentially be explained by spurious sequencing of RNA transcripts, potentially through reverse transcription during library preparation. To investigate this hypothesis, we posit that in the absence of sequencing artefacts, 5’ and 3’ paired-end metagenomic reads should be similarly mapped upstream and downstream of the TSS. However, in the presence of RNA fragment sequences, we would expect an enrichment of 5’ reads immediately downstream the TSS compared to 3’ reads (**Fig. 4a**).

**Figure 4.**
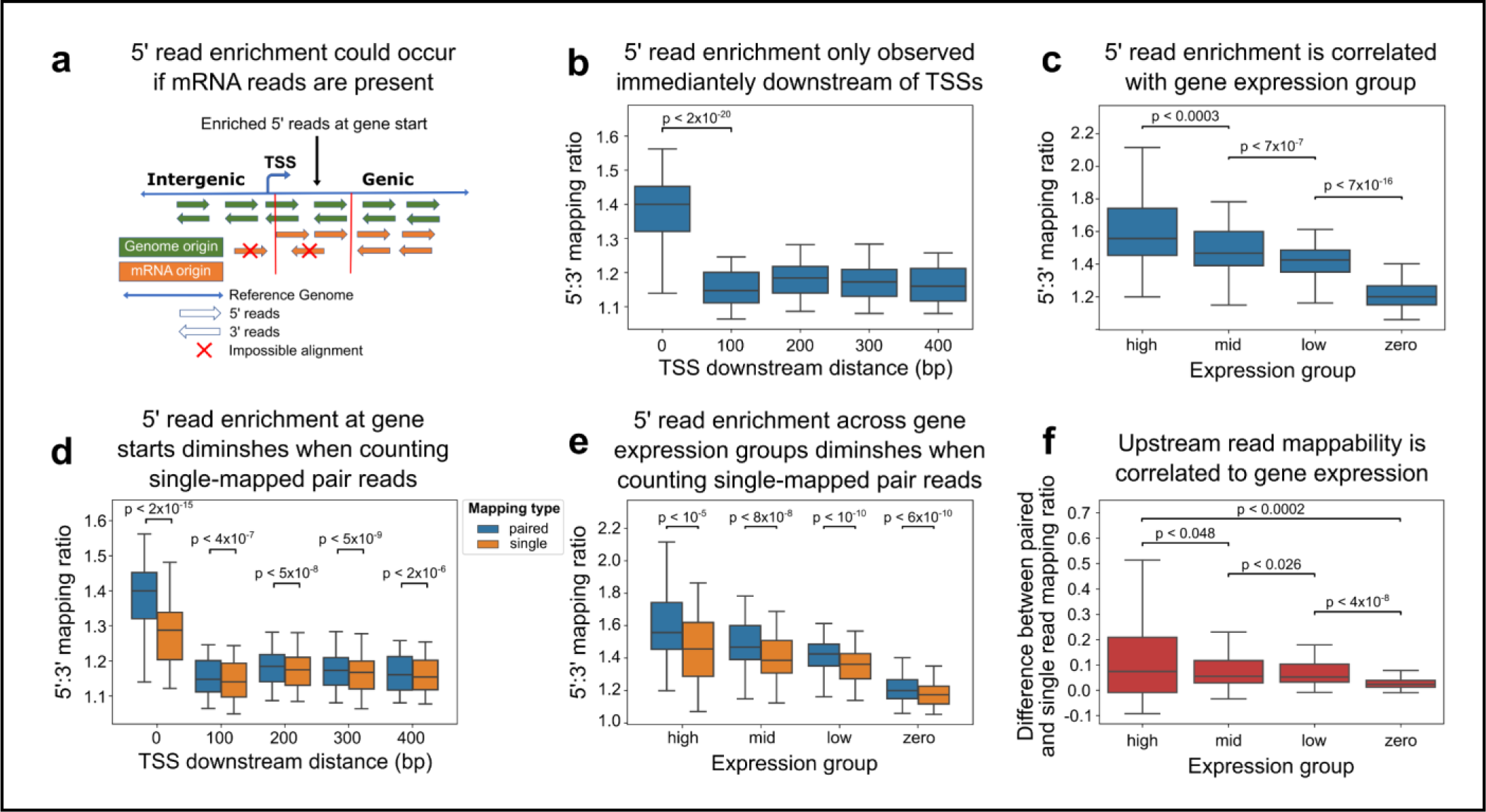
Mappability bias upstream of TSS is associated with gene expression. **(a)** Diagram illustrating a potential effect of spurious RNA sequencing on TSS coverage bias. 5’ read enrichment downstream of TSSs suggests the presence of sequenced mRNA. **(b)** Boxplot (line, median; box, IQR; whiskers, 1.5xIQR) of 5’:3’ mapping ratio at different 50bp windows downstream the TSS. The 5’:3’ mapping ratio is calculated as the ratio between total 5’ reads and 3’ reads in a given window, where only reads with both ends mapped are considered. **(c)** Boxplot of 5’:3’ mapping ratio for the window immediately downstream of the TSS, across different gene expression groups. **(d)** Boxplot of 5’:3’ mapping ratio at different 50bp windows of the gene starting from the TSS, using two methods for read counting. “Paired” only considered reads in which both ends were successfully aligned, whereas “single” also considered reads where only one end was successfully aligned. **(e)** Boxplot of 5’:3’ mapping ratio across different gene expression groups, using two methods for read counting (“paired” and “single”). **(f)** Difference in 5’ read enrichment when performing “paired” and “single” read counting, across different expression groups. For all analysis, datapoints represented the averaged value of a sample and species, where values from reference genomes of the same species were averaged. Statistical significance across groups was determined under the paired Student’s t-test where values were paired by their sample and species.

We therefore analysed the distribution of paired-end read mappings in all 8 samples from the Franzosa et al. dataset analysed above (Methods). We indeed found an enrichment of 5’ reads compared to 3’ reads in the 50 bp window immediately downstream of the TSS, with 1.4x (1.39±0.087) more 5’ reads (**Fig. 4b**). This was significantly reduced 100bps downstream of the TSS (paired t-test, p = 1.45x10^-20^, **Fig. 4b**), with only 1.15x (1.15±0.054) more 5’ reads. Although it is uncertain why there is this minor enrichment in 5’ reads throughout the gene, the dramatic enrichment in 5’ prime reads at gene starts supports our hypothesis that RNA fragments may be contributing to coverage. To validate these results we checked whether 5’ mapping difference was associated with gene expression, where highly expressed genes are expected to have more RNA and therefore more 5’ read coverage. We found 5’ read enrichment indeed positively correlated with gene expression group (Kruskal p = 5.90х10^-22^, **Fig. 4c**).

We further interrogated whether this difference in 5’ and 3’ read coverage at the gene start is due to read mapping biases between genic vs. intergenic regions. If the 5’ pair originated upstream of the TSS, in an intergenic region, and fails to map due to mappability, it will cause a reduction in 3’ reads downstream of the TSS. We therefore repeated the same analysis as above, but counted read pairs in which only one of the reads mapped (“single-mapped read pairs”). With the inclusion of such read pairs, we found that the ratio of 5’ read enrichment decreased significantly to 1.28x (1.28±0.091) (paired t-test p = 1.47х10^-17^, **Fig. 4d**), however this was still higher than 5’ read enrichment 100 bps downstream of the gene (paired t-test p = 8.42x10^-12^, **Suppl. Fig. 4a**), which had an average 5’ read enrichment of 1.14x (1.14±0.055). The decrease in 5’ read enrichment at the gene start after counting single-mapped read pairs strongly indicates a mappability bias upstream of the TSS. This however does not entirely explain the total 5’ read enrichment at the gene start, which is still significantly higher compared to regions downstream of the gene (**Suppl. Fig. 4a**). This residual effect could potentially suggest sequencing of RNA.

When investigating the effects of incorporating single-mapped read pair counting to the correlation with gene expression groups, we see a significant decrease in the 5’:3’ mapping ratio in all expression groups (**Fig. 4e**). There still persists a positive correlation between 5’ read enrichment with expression group, although weaker than previously (Kruskal p = 6.51х10^-19^, **Suppl. Fig. 4b**). We next measured the difference in 5’ read enrichment between paired read and single-mapped read counting, and demonstrated highly expressed genes tend to have a greater decrease in 5’ read enrichment after including single-mapped reads compared to lowly expressed genes (Kruskal p = 0.00084, **Fig. 4f**). To validate this was not an artefact of gene dependency across species/samples, we modelled this association in a linear mixed model allowing sample and species information to be included as random effects. This showed gene expression to be positively associated with the difference in paired and single-mapped read counting (p < 1.05x10^-05^, **Suppl. Table 1**). This indicates that the degree of mappability bias occurring upstream of the TSS is correlated with the expression of the gene.

### TSS coverage bias is absent when aligning to strain-specific reference

To further investigate the hypothesis that 5’ reads upstream of the TSS are unmapped due to strain differences between the sample-specific genomes and the reference species genomes used for alignment, we performed the same analysis on metagenomic sequencing of a 20-member mock community (28), for which the exact reference genome of all species is known (Methods). We found no 5’ read enrichment downstream of the TSS, regardless of whether single-mapped read pairs were considered (**Fig. 5a**). We further show the absence of average coverage bias at TSSs across all species in the mock community (**Suppl. Fig. 5**). This result supports a hypothesis of 5’ read enrichment generated due to strain heterogeneity.

**Figure 5.**
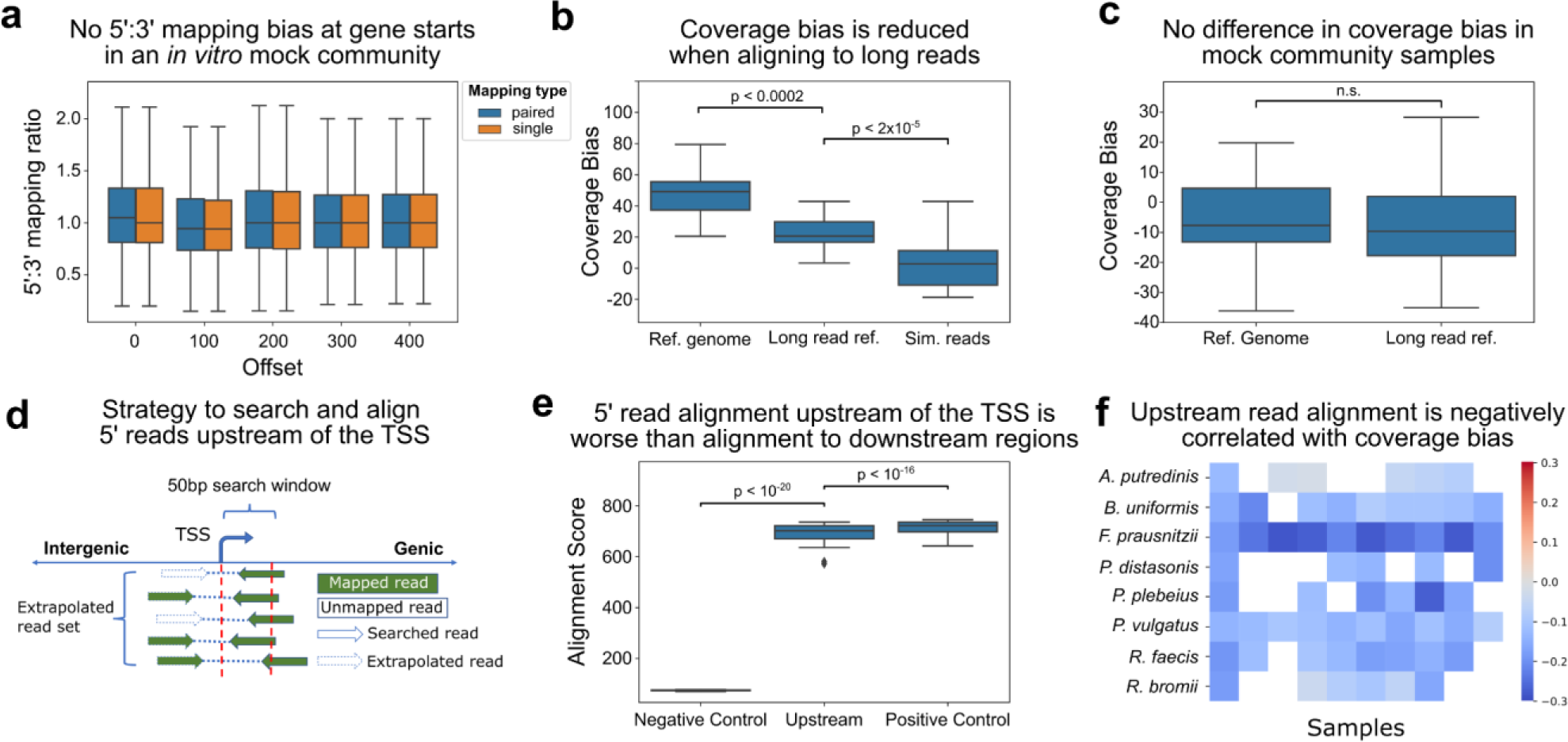
Sequence heterogeneity upstream of the TSS contributes to coverage bias. **(a)** Boxplot (line, median; box, IQR; whiskers, 1.5xIQR) of 5’:3’ mapping ratio at different 50bp windows of the gene starting from the TSS for an *in vitro* mock community sample. Reads were aligned to all known species genomes (N = 20) in the community. **(b)** Boxplot of coverage bias from 10 gut metagenome samples when aligning short reads to reference genome, aligning the same short reads to reference generated from matched long reads, and aligning simulated short reads to the same long read reference. The average bias across species genomes was plotted for each sample. **(c)** Boxplot of coverage bias from a mock community of 20 bacterial species when aligning short reads to reference genome and matched long read reference. **(d)** Diagram demonstrating the strategy to search and align 5’ reads upstream of the TSS to the reference genome. All 3’ reads mapped immediately downstream of the TSS are first identified. Their 5’ paired read is then aligned to the reference genome via global cost-free ends alignment. **(e)** Boxplot of 5’ read alignment scores under three frameworks. Negative control aligns randomly selected reads to the 300 bp upstream to 200 bp downstream of the TSS. “Upstream” denotes alignment of 5’ reads whose 3’ pair mapped downstream of the TSS (as described in panel d). Positive control captures the alignment of 5’ reads downstream of the TSS. This was performed across 10 gut metagenome samples and selected species genomes. Alignment scores for each TSS were averaged for each sample and reference genome. **(f)** Boxplot of Pearson R score between upstream alignment score and coverage bias correlation across species. Correlation and scores for reference genomes of the same species were averaged. P - paired t-test.

To further investigate whether TSS coverage bias is due to strain heterogeneity, we next analysed a dataset in which paired Illumina short-read sequencing and Oxford Nanopore long-read sequencing was performed on 10 stool samples from healthy individuals (25). We then mapped short reads to long reads from the same sample. This alleviates bias caused by inter-sample strain diversity (i.e., difference between strain in the samples to the reference genome), but not bias caused by intra-sample strain diversity (i.e., differences between strains within the same sample). We considered all species with at least 1% abundance in at least 5 samples, and compared the coverage bias when (1) mapping short reads to reference genomes from Refseq following the same method for coverage bias calculation as before (2) mapping to long-reads from the same sample, and calculating the average TSS coverage bias per reference genome around genes predicted with Prodigal (29) (Methods). Indeed, we observed a significant reduction in coverage bias when aligning to long-reads compared to when aligning to reference genomes (23.26 vs. 45.6, respectively; paired t-test p=0.00016, **Fig. 5b**). We note, however, that the TSS coverage bias was still present, even when mapping to long-reads, potentially reflecting the effect of strain diversity within a sample. A negative experiment of aligning simulated short-reads to long reads had significantly lower bias at TSSs (paired t-test, p = 1.82x10^-5^, **Fig. 5b**).

To exclude the possibility of an artefact arising from the segmented nature of long reads used as a reference, we repeated this experiment in the previously mentioned bacterial mock community sample, for which PacBio Hi-Fi long read sequencing data was available. For this sample, exact reference genomes are available, and intra-sample strain diversity should be minimal. We found no significant difference between coverage bias when aligning to the reference genome versus long read references (paired t-test p = 0.77, **Fig. 5c**).

To directly investigate the hypothesis of strain diversity causing a bias upstream of the TSS, we investigated the alignment quality of reads from ten stool samples from the ARG dataset (23) for abundant taxa species (at least 1% abundance in at least 5 samples). This dataset contained longer fragment lengths allowing the distance of paired reads to span across the TSS with minimal dovetailing. To this end, we aligned the 5’ ends of all read pairs whose 3’ end mapped to the 50 bps downstream of the TSS to the 500 bp region upstream of the TSS using global cost-free-ends alignment (**Fig. 5d**, Methods). The alignment quality of these reads serves as an indication for strain diversity upstream of the TSS, where lower qualities indicate higher strain diversity. We also performed two controls. As a “negative control”, to ensure that lower alignment scores are not due to spurious misaligned reads, we aligned randomly selected 5’ read ends to the same region. As a positive control, we performed the same analysis, only instead of using the regions up/downstream of the TSS, we selected regions that are downstream of the TSS (using the average fragment length of the sample), such that both read ends are mapped downstream of the TSS. Indeed, we found that the average (across samplexspecies) alignment quality of 5’ read ends upstream of the TSS was significantly higher than the alignment quality in the negative control (paired t-test p < 10^-20^), and significantly lower than those of the positive control (paired t-test p = 1.42х10^-16^, **Fig. 5e**). Furthermore, we demonstrated the 5’ read alignment quality was negatively associated with coverage bias in a linear mixed model with sample and reference genome as random effects (p < 2x10^-16^, **Suppl. Table 2**). When looking at the Pearson correlation for individual species, we observe significant associations for all species except *Ruminococcus bromii* and *Alistipes putredinis* (**Fig. 5f, Suppl. Fig. 6**). Altogether, these analyses strongly suggest that coverage bias is caused by strain diversity.

## Discussion

In this study we demonstrate the value of investigating sequencing coverage patterns in metagenomic data, by detecting a phenomenon of coverage bias at transcription start sites which coincides with gene expression. We validated this phenomenon systematically across datasets and demonstrated that it exists across microbial species with sufficient coverage. Extensive investigation of potential artefacts ruled out previously known technical artefacts such as read mappability bias and sequencing platform biases. Although bias in coverage was correlated with GC bias, its association with gene expression was independent of this relationship. By examining paired-end reads that align immediately downstream of the TSS, we found there was an enrichment of 5’ compared to 3’ reads, which could indicate sequencing of RNA transcript sequencing. However, this 5’ read enrichment diminished when also considering reads pairs in which only one pair was successfully mapped. We hypothesise this was due to strain sequence variation from reference genomes. We first validated the absence of 5’ read enrichment in a mock community of 20 bacterial species where the exact strain reference genome is known. Additionally, we showed that TSS coverage bias significantly decreased when aligning short-reads to matched long-reads whereas TSS coverage bias did not exist in *in vitro* mock samples regardless of alignment to reference genome or long-read reference. Sequence homology of reads upstream of the TSS proved to be more divergent to the reference genome used for alignment compared to reads downstream of the TSS. The divergence of upstream reads was also significantly correlated with coverage bias, indicating a major source for coverage bias at the TSS is due to alignment biases as a result of unique community strains.

We did not observe the complete abolishment of 5’ read enrichment (i.e. a ratio of 1) after accounting for unmapped read pairs. This suggests that unintended RNA transcript sequencing may still be a plausible phenomenon, although the complete lack of TSS coverage bias in mock communities is strong evidence against it. The degree of 5’ read enrichment correlated with gene expression, where highly expressed genes had a greater reduction in 5’:3’ read mapping ratio when considering unmapped read pairs. This indicates that heterogeneity-derived alignment bias upstream of the TSS is positively correlated with gene expression, where highly expressed genes would contain more divergent promoter sites as reflected by greater coverage bias. Although this correlation is not casual or directional, we propose a plausible hypothesis that mutational signatures in the promoter/enhancer region are required for the regulation of gene expression in a context-specific manner.

Due to the highly competitive and changing environment of the gut microbiome (30), strong evolutionary pressures exist, which can induce specific mutational imprints on bacterial genomes (31). This likely affects sequence divergence of promoter sites and optimised gene expression (32). The mechanism for this was demonstrated in studies of the prokaryotic promoter sites where random sequences were shown to rapidly evolve towards functional promoters (33). The inclusion of mutational signatures within promoter sites were additionally demonstrated to improve biophysical models to predict gene expression levels (34). An evolutionary study in *E. coli* demonstrated the build- up of mutations in cis-regulatory regions induced changes to the intrinsic properties of existing promoters as well as produced new transcription sites (35). The malleability of promoter sites for gene expression and flexibility of polymerase binding would support the idea of evolutionary finetuning of gene expression through promoter site mutation. In addition, there is support for the evolutionary dynamics of gut microbiota to occur within human-relevant timescales, where both short-term and long-term evolution exists with hosts (36, 37).Collectively, we foresee this evidence to be very applicable to the gut microbiome where its vast functional complexity and species diversity likely arises from evolutionary adaptation to environmental stresses of the gut. The positive association observed in our study between gene expression and promoter heterogeneity perhaps reflects the importance of regulatory control on highly expressed genes and thus justifies increased evolutionary footprints within their promoter sequences. Future longitudinal experiments in *in vivo* systems that capture the evolutionary forces applied on gut microbial genomes would be helpful to further validate this hypothesis.

We acknowledge that this study is limited to bioinformatic analysis utilising public datasets originally produced with different research aims. This limits us from performing tailored perturbation experiments, with the potential to test alternative biological hypotheses such as supercoiling and promoter binding influencing coverage and mutation rates at promoters. This study would also have greatly benefited from the existence of culturomics data of strain genomes from gut microbiomes for the validation of divergence occurring at intergenic promoter sites. It would also be valuable extending our hypothesis and analysis to other complex environments with high ecological pressure such as the soil microbiome to investigate reproducibility. Nevertheless, our work illuminates the potential for evolutionary footprints of strain adaptation within gut metagenomes which is revealed as a byproduct of biases in coverage due to divergence from reference genomes. This empowers our understanding of metagenomic read data, its potentially adverse effects on downstream methods that assume even coverage, as well as possible strain-level evolutionary behaviour that may manifest from different ecology pressure across individual gut microbiomes. We hope this paves an avenue of future work elucidating mechanisms of gene regulation through metagenomic data.

## Data availability

All raw sequencing data is publicly available on SRA as described in the Methods section.

## Code availability

Code to replicate all analysis steps is available from https://github.com/holab-hku/TSS_project.

## Acknowledgments

We thank members of the Korem and Ho groups for useful discussions. This work was supported in part by AIR@InnoHK administered by Innovation and Technology Commission of Hong Kong, the Hong Kong PhD Fellowship Scheme of the Research Grants Council of Hong Kong, the Bau Tsu Zung Bau Kwan Yeu Hing Research and Clinical Fellowship, the Program for Mathematical Genomics at Columbia University, and the CIFAR Azrieli Global Scholarship in the Humans & the Microbiome Program.

## Author contributions

Gordon Qian - Conceptualization, Formal analysis, Methodology, Investigation, Visualisation, Writing – original draft, Writing – review and editing Izaak Coleman - Methodology, Investigation Tal Korem - Conceptualization, Methodology, Investigation, Writing – review and editing, Supervision Joshua Ho - Conceptualization, Methodology, Investigation, Writing – review and editing, Supervision

## Supplementary figures

**Supplementary Figure 1.**
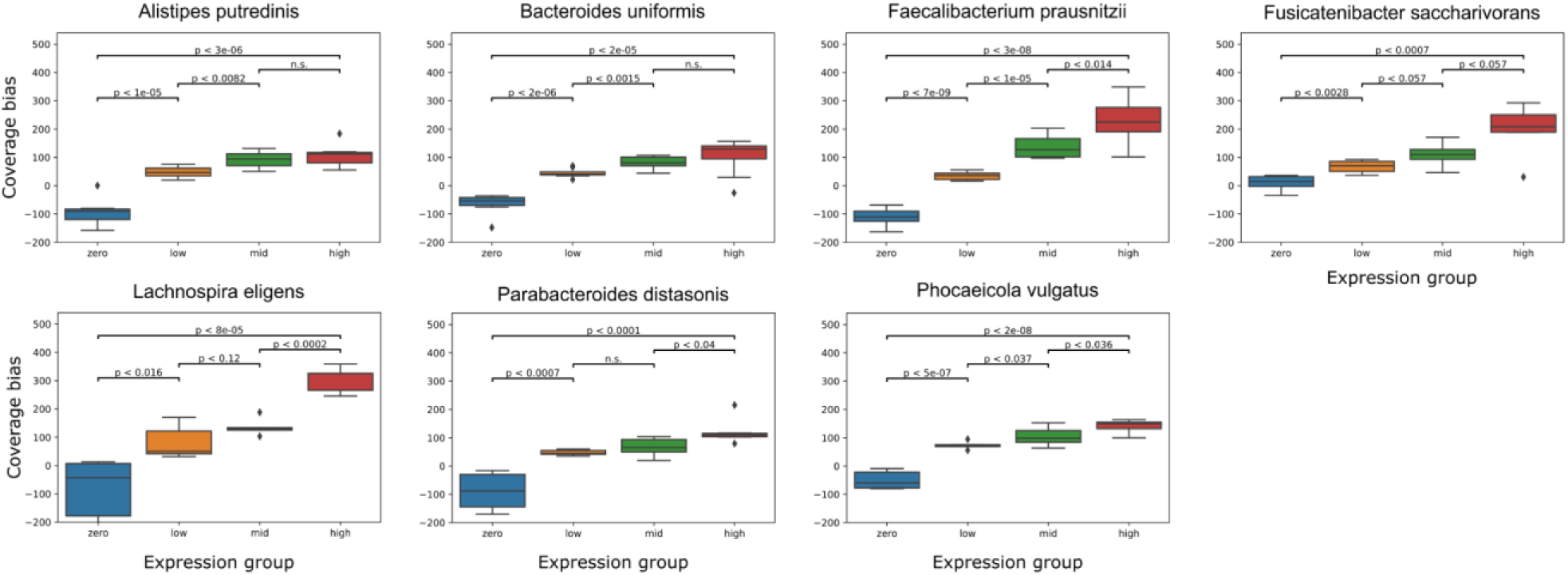
Species-stratified analysis of coverage bias and gene expression groups. Boxplot of TSSs coverage bias against gene expression groups for each individual species. Statistical significance between groups was performed under the paired Student’s t-test. P value of < 0.05 was considered significant.

**Supplementary Figure 2.**
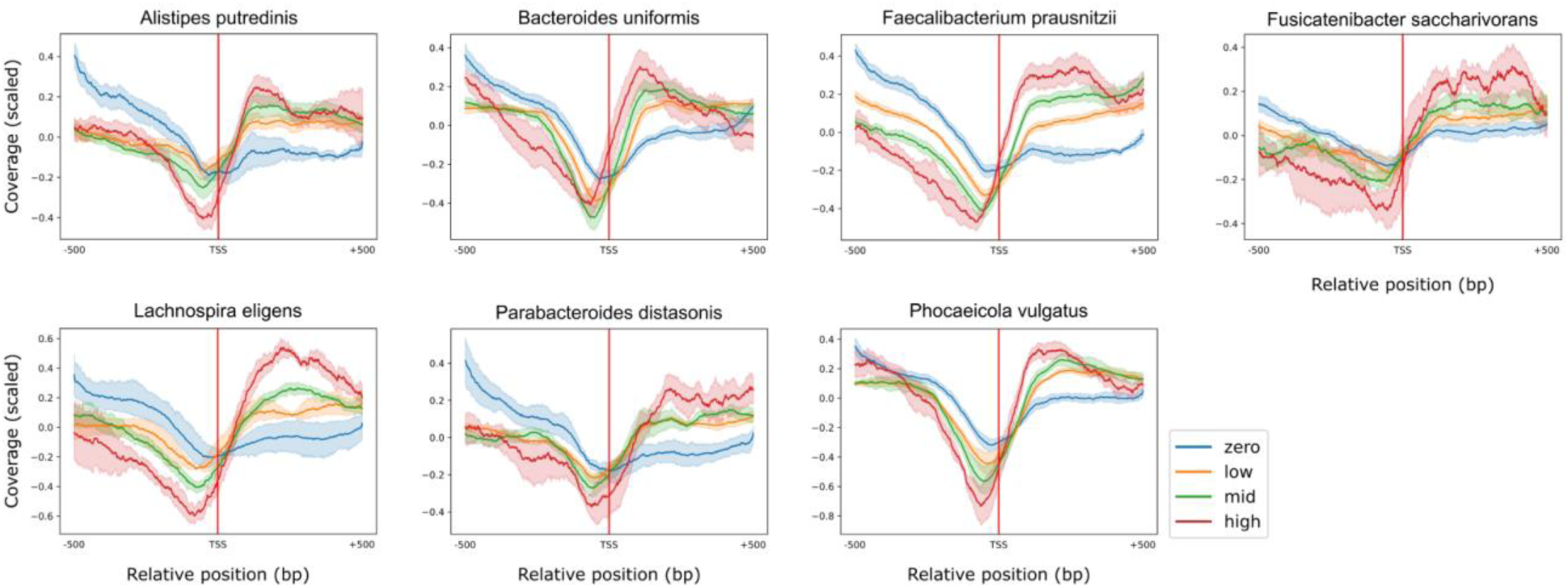
Species-stratified scaled coverage across TSS locus. Line plot of scaled coverage across the 1 kbp TSS window where TSSs are grouped by gene expression for each species. Coverage values averaged at each individual base pair position in the window. Measure of spread represents the 95% CI.

**Supplementary Figure 3.**
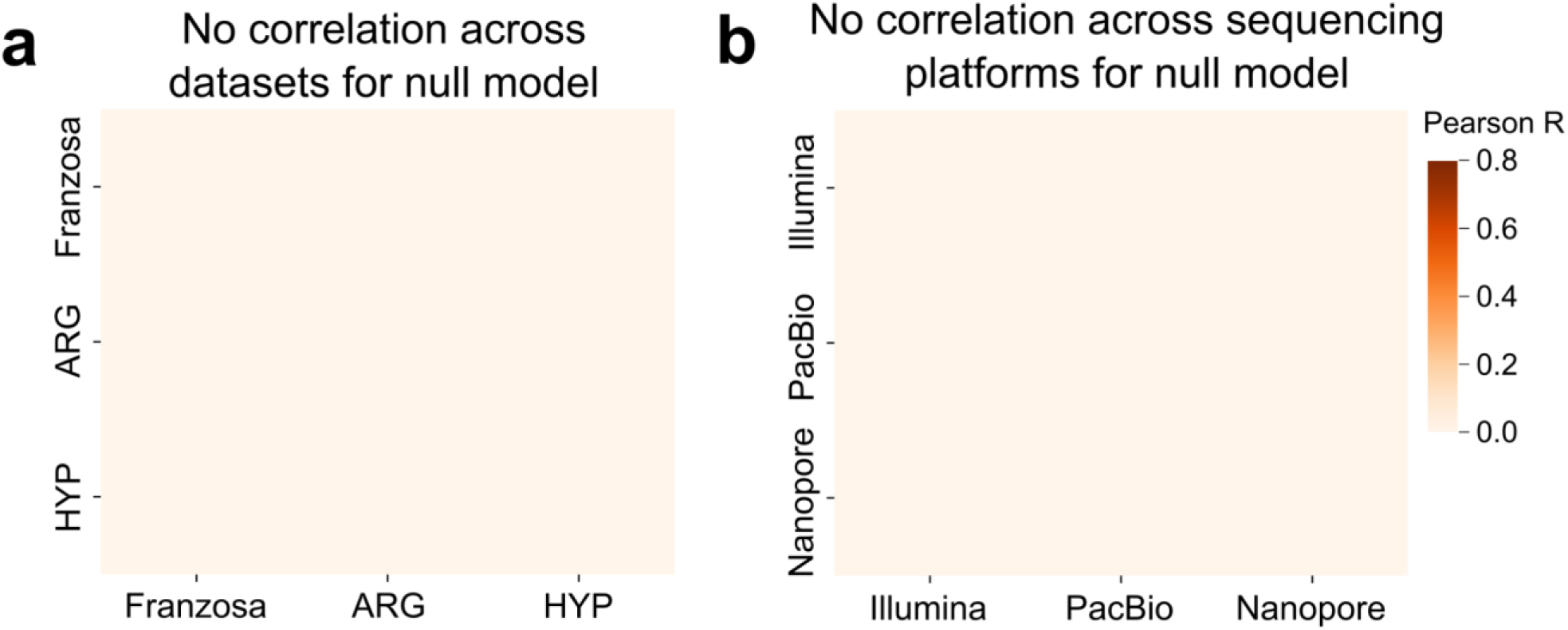
Negative controls for coverage correlation. Heatmap of Pearson correlation scores between pairwise sample coverage bias from different **(a)** Illumina short-read datasets and **(b)** sequencing platforms, that were calculated after randomly permuting coverage.

**Supplementary Figure 4.**
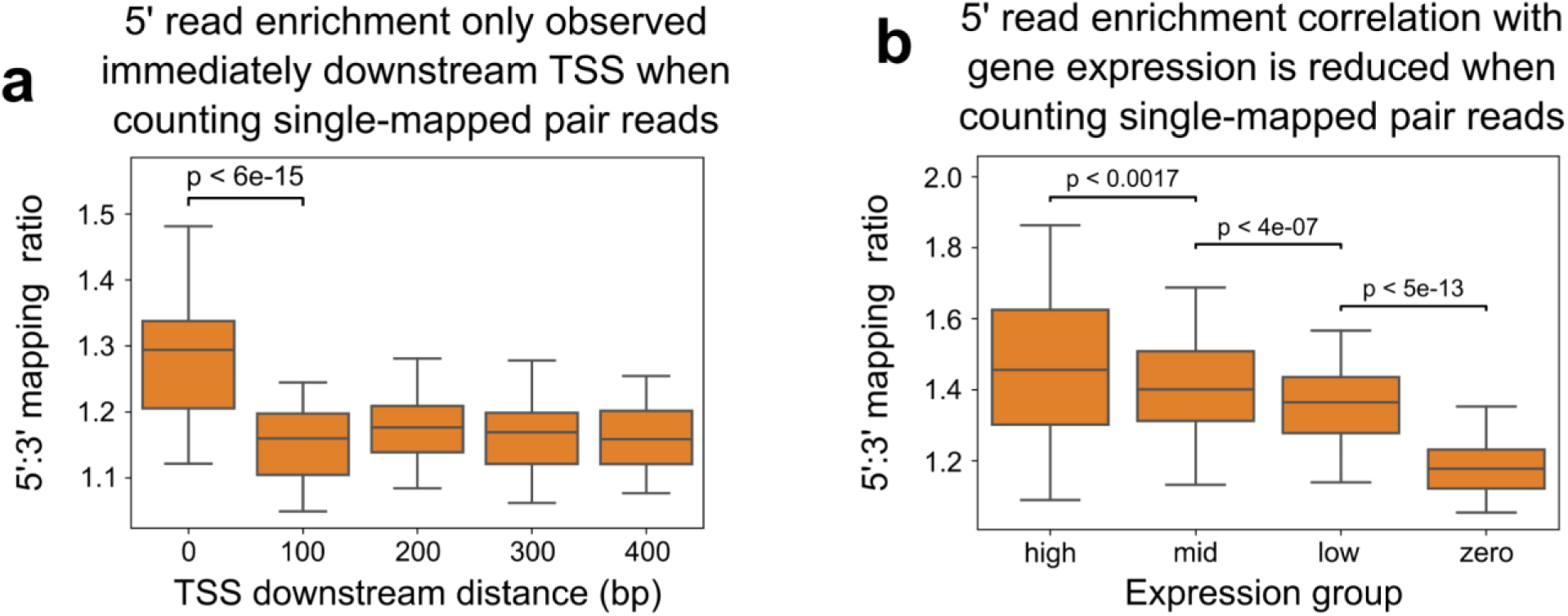
5’ read enrichment analysis with “single” paired-end read mapping counting. Boxplot of 5’:3’ mapping ratio (a) at different 50bp windows of the gene starting from the TSS, (b) across gene expression groups, under “single” paired-end read counting. Datapoints represented the averaged value of a sample and species, where values from reference genomes of the same species were averaged. Statistical significance across groups was determined under the paired Student’s t-test where values were paired by their sample and species.

**Supplementary Figure 5.**
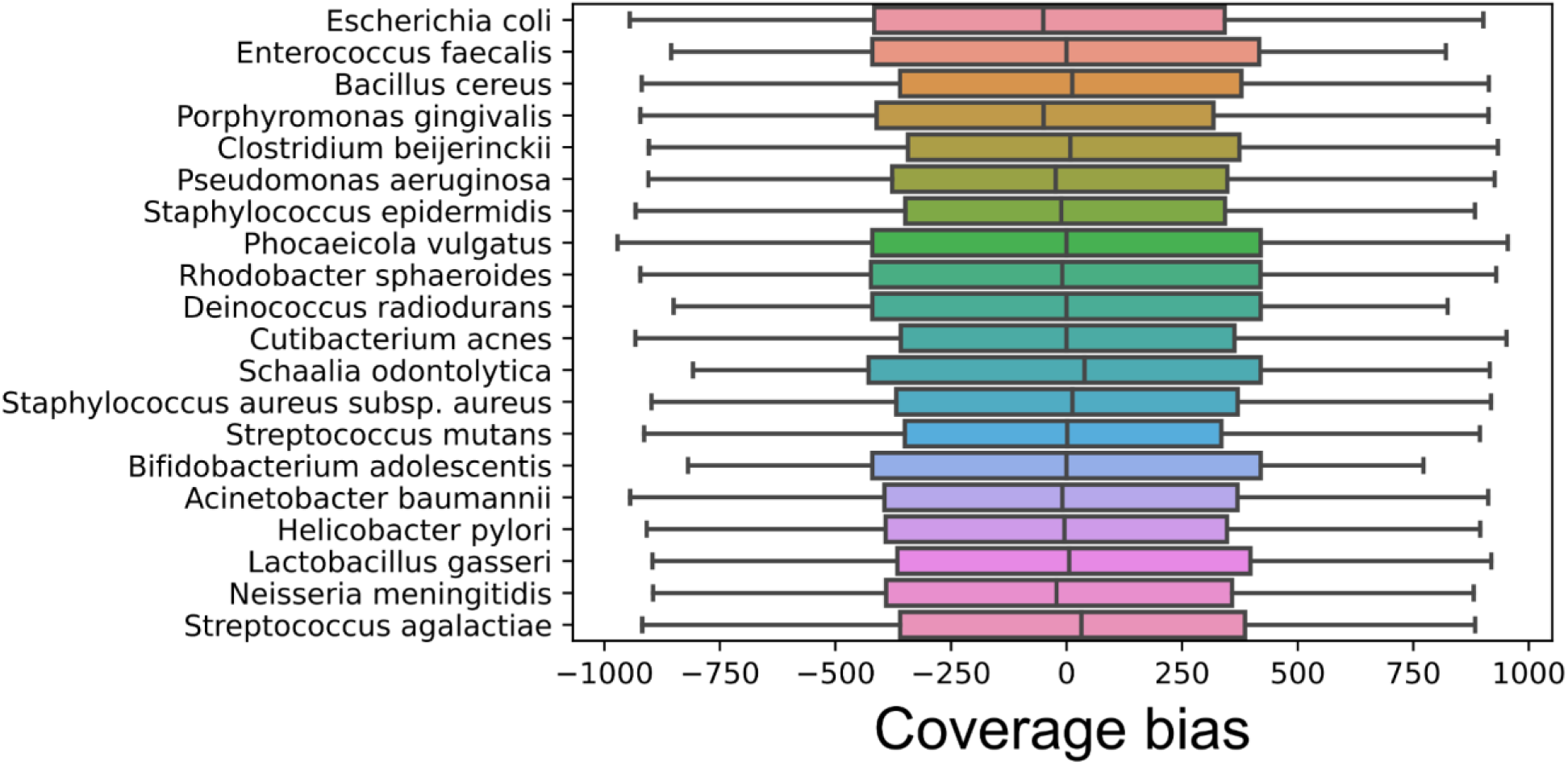
Coverage bias at TSSs do not exist in species from an *in vitro* mock community. Boxplot (line, median; box, IQR; whiskers, 1.5xIQR) of coverage bias for each species in a 20-member mock community when aligning short reads to their exact reference genome.

**Supplementary Figure 6.**
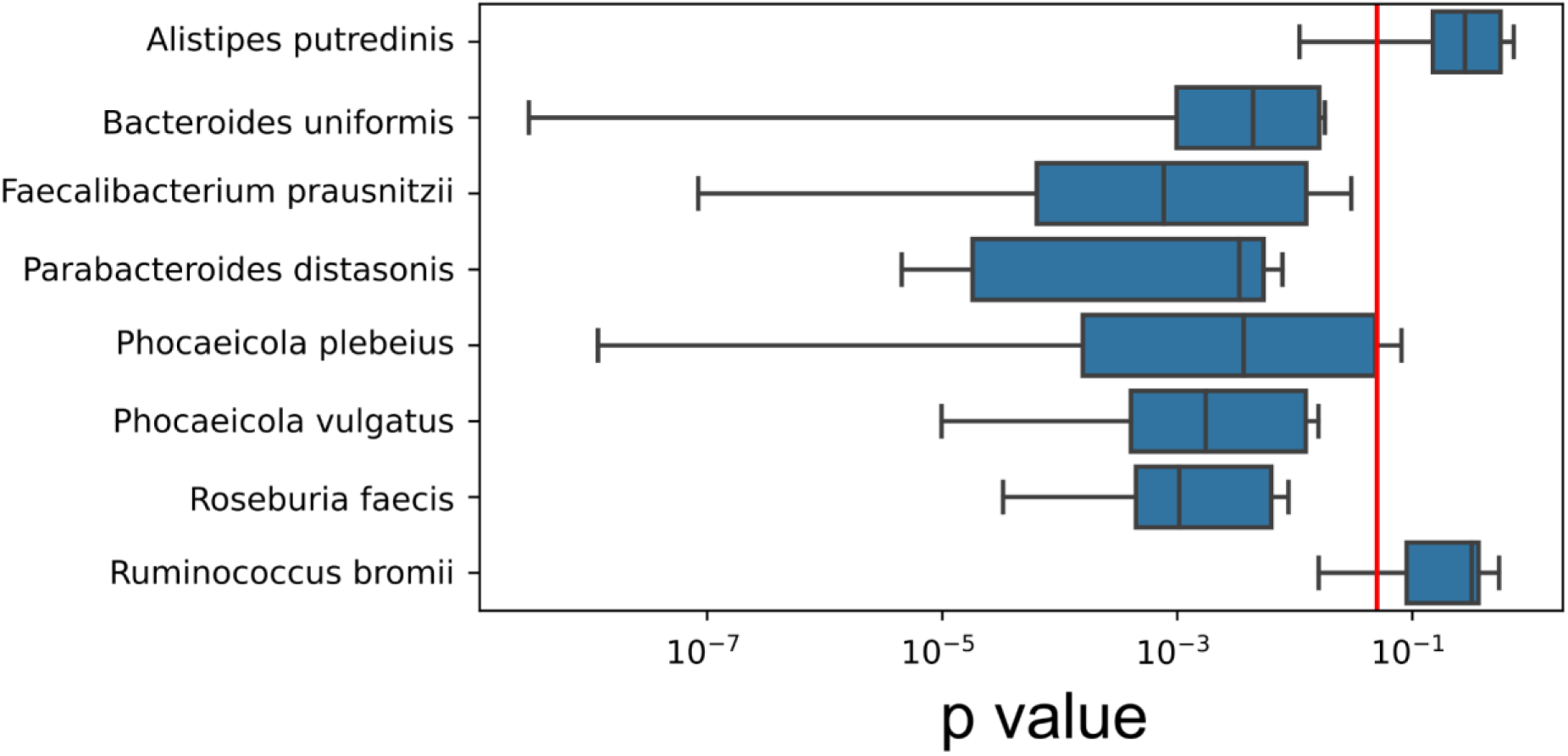
Coverage bias is significantly associated with the alignment of reads upstream to the TSS. Boxplot (line, median; box, IQR; whiskers, 1.5xIQR) of p-value scores from the Pearson correlation test between upstream alignment score and coverage bias across different species from the ARG dataset. The vertical red line indicates the threshold for significance of p = 0.05.

## Supplementary tables

**Suppl. Table 1.** Gene expression is positively associated with the difference in 5’ read enrichment between “paired” and ”single” read counting.

| Linear Mixed Model | Estimate | Standard Error | t value | P > \|z\| |
| --- | --- | --- | --- | --- |
| 5' read enrichment difference between paired and single-mapped read counting | $3.61 \times 10^{-2}$ | $1.61 \times 10^{-3}$ | 22.39 | $< 2 \times 10^{-16}$ |

**Legend.** Gene expression was modelled under a linear mixed model with the fixed effect, the difference in 5':3' mapping ratio between "paired" and "single" read counting. Random effects, sample and species, allowed grouping of observations. Gene expression (TPM) values were log-transformed, scaled and centered. 5' read enrichment variable was also scaled and centered. There were a total of 354,697 observations, 20 species groups and 8 sample groups. Model convergence by Restricted Maximum Likelihood (REML).

**Suppl. Table 2.** Coverage bias is negatively associated with the alignment score of unmapped reads upstream of the TSS.

| Linear Mixed Model | Estimate | Standard Error | t value | P > \|z\| |
| --- | --- | --- | --- | --- |
| Upstream alignment score | $-1.55 \times 10^{-1}$ | $3.22 \times 10^{-3}$ | -48.11 | $< 2 \times 10^{-16}$ |

**Legend.** TSS coverage bias was modelled under a linear mixed model with the fixed effect, upstream read alignment score. Random effects, sample and species, allowed grouping of observations. Coverage bias and alignment score variables were scaled and centered. There were a total of 101,120 observations, 24 species groups and 10 sample groups. Model convergence by Restricted Maximum Likelihood (REML).

